# Endogenous SIRT3 activity is dispensable for normal hearing recovery after noise exposure in young adult mice

**DOI:** 10.1101/2020.02.05.935882

**Authors:** Sally Patel, Lisa Shah, Natalie Dang, Xiaodong Tan, Anthony Almudevar, Patricia M. White

## Abstract

Occupational noise-induced hearing loss (NIHL) affects millions of people worldwide and presents a large social and personal burden. Some genetic variants in the mitochondrial oxidative stress response correlate strongly with susceptibility to NIHL in both humans and mice. Here we test the hypothesis that SIRT3, a regulator of the mitochondrial oxidative stress response, is required in mice for endogenous recovery of auditory thresholds after a sub-traumatic noise exposure. We expose homozygous *Sirt3-KO* mice and their wild-type littermates to a noise dose that confers a temporary threshold shift, but is not sufficient to permanently reduce cochlear function, compromise cell survival, or damage synaptic structures. We find no difference in hearing function after recovery from noise exposure between the two genotypes, when measured by either auditory brainstem response (ABR) or distortion product otoacoustic emissions (DPOAE). We show that SIRT3-specific immunoreactivity is present in outer hair cells, around the post-synaptic regions of inner hair cells, and faintly within inner hair cells. Nonetheless, outer hair cells and auditory synapses show no increase in loss after noise exposure in the homozygous *Sirt3-KO* mouse. These data show that SIRT3-dependent processes are not necessary for endogenous hearing recovery after a single, sub-traumatic noise exposure. They demonstrate the existence of cellular mechanisms of cochlear homeostasis in addition to the mitochondrial oxidative stress response. We also present a novel statistical analysis for identifying differences between peak 1 amplitude progressions in ABR waveforms.

## INTRODUCTION

Noise-induced hearing loss (NIHL) affects at least 10 million adults in the United States [1], including over a million veterans [2]. Noise-induced auditory dysfunction, including tinnitus and NIHL, is the most common disability among former combat soldiers, costing the Veteran’s Administration over a billion dollars annually [2]. NIHL is a form of acquired hearing loss, which can be associated with greater levels of anxiety [3], emotional distress [4], and perceived stigmatization [5], as well as poorer health outcomes [6]. There is no approved biological treatment for NIHL [1], and it can only be prevented by physically avoiding noise exposure.

Recent progress has been made in identifying genetic variants that predispose individuals to NIHL, especially in occupational settings [7, 8]. Gene variants encoding proteins that modulate oxidative stress are over-represented in these studies (for review, see [9]). More specifically, gene variants that reduce mitochondrial function also enhance susceptibility to acquired hearing loss from noise [10, 11], ototoxic drugs [12], and age-related hearing loss [13, 14]. These facts support the interpretation that proteins promoting mitochondrial function and counteracting oxidative stress also protect the cochlea from noise damage. SIRT3 is a mitochondrial lysine deacetylase [15] that promotes an effective oxidative stress response [16] from mitochondrial enzymes, including Superoxide dismutase 2 (SOD2, [17]). SIRT3 variants are also implicated in human longevity [18]. SIRT3 activation has previously been shown to protect cochlear outer hair cells (OHCs) from aging, where investigators used dietary restriction to activate sirtuins. The positive effect was only observed in wild-type, not homozygous *Sirt3-KO* mice [19]. Importantly, genetic or pharmaceutical activation of sirtuins was also shown to rescue hearing from traumatic noise damage in a SIRT3-dependent manner [20].

Based on its function and these previous SIRT3 gain-of-function findings, we investigated whether there is a requirement for endogenous SIRT3 in the normal recovery of hearing thresholds in adult mice after a temporary threshold shift (TTS). Since TTS is associated with a recoverable disrupted cellular process rather than hair cell loss [21], the role of SIRT3 for restoring mitochondrial function and hearing recovery in subtraumatic noise could be determined by comparing wild-type and *Sirt3-KO* mice. These loss-of-function studies also complement the already-published gain-of-function studies. Multiple labs have successfully used this method to identify genes [22] and conditions [23] that modulate susceptibility to noise damage. Noise exposure can eliminate high-frequency auditory synapses [21, 24] in a glutamine-dependent manner [25], and the restoration of auditory synapses was proposed to be the mechanism by which SIRT3 reduced NIHL [20]. To evaluate any requirement for endogenous SIRT3 activity in resilience from noise damage, we employed a noise exposure that is 80% of the energy level needed to induce synaptopathy [21, 24]. It induces small or negligible permanent ABR threshold shifts [22], and does not cause OHC death [22]. By exposing homozygous *Sirt3-KO* mice and their wild-type littermates to this subtraumatic noise, we sought to uncover a role for endogenous SIRT3 activation in preserving these cellular structures and functions.

## MATERIALS AND METHODS

### Animal usage

All experiments were performed in compliance with the US Department of Health and Human Services, and were approved by the University Committee on Animal Resources at the University of Rochester Medical Center or the IACUC at Northwestern University. Male *Sirt3-KO* mice (129-Sirt3tm1.1Fwa/J; stock number 012755; The Jackson Laboratory) were bred four times to FVB/nJ females (stock number 001800, The Jackson Laboratory). In other experiments, we have found that four generations is sufficient to confer youthful noise damage sensitivity similar to that of congenic FVB/nJ [22, 24]. Heterozygotes with different parents were bred together to obtain both knockout and wild-type littermates. Both males and females were used. In the first five litters generated, fourteen homozygous *Sirt3-KO* and thirteen wild-type littermates were generated and then exposed to noise. Four additional litters were generated to produce homozygous *Sirt3-KO* mice and wild-type littermates, which were not exposed to noise, for histology. Mice were given ample nesting materials and small houses within their home cage.

For genotyping, DNA was obtained from 2-mm tail samples that were digested overnight in Proteinase K (IBI Sciences) solution at 65°C followed by phenol/chloroform extraction. The KAPA Taq PCR kit (Sigma, BK1000) was used in conjunction with three primer sequences (wild type, CTT CTG CGG CTC TAT ACA CAG; common, TGC AAC AAG GCT TTA TCT TCC; mutant, TAC TGA ATA TCA GTG GGA ACG) to identify genotypes.

### Noise exposure

Awake two month old mice were exposed to noise limited to the 8-16 kHz octave band at 105 decibels for 30 minutes. Mice were each placed into individual triangular wire mesh cages, 12 cm x 5 cm x 5 cm, in an asymmetric plywood box with a JBL2250HJ compression speaker and JBL2382A biradial horn mounted on the top. This apparatus was contained within a sound booth. The speaker was driven by a TDT RX6 multifunction processor and dedicated attenuator, and controlled with TDT RPvdsEx sound processing software. The sound level was calibrated with a Qwest sound meter, and the sound level was checked each morning with a calibrated iPhone using the FaberAcoustical SoundMeter app. Mice were exposed to noise between 9 am and 4 pm.

### Auditory testing

Mice were tested at 7 weeks of age (pre-test), one day after receiving noise exposure, and again fourteen days later. Mice were exposed to noise at P60. Auditory testing was conducted using a Smart EP Universal Smart Box (Intelligent Hearing Systems) with high-frequency speakers from Tucker Davis Systems. Mice were anesthetized with an intraperitoneal injection of ketamine (80 mg/kg) in a sterile acepromazine/saline mixture (3 mg/kg). A 10B+ (high frequency transducer/stimulator) probe was placed at the opening to the external auditory meatus. Sound production in this system was calibrated with a Qwest sound meter every six months.

Auditory brainstem response (ABR) stimuli were 5-ms clicks, or 5-ms tone pips presented at 5 frequencies between 8 and 32 kHz. Stimuli began at 75 dB amplitude and decreased by 5 dB steps to 15-25 dB. 512 sweeps were averaged for each frequency and amplitude. Electrical responses were measured with three subdermal needle electrodes (Grass): one inserted beneath each pinna, and a third, the ground electrode, placed at the vertex. ABR thresholds for a particular frequency were determined by any part of the last visible trace (dB). The person scoring the waveforms was blinded to genotype and time point. Masking and randomization was performed by drawing cards from a fair deck and renaming files appropriately.

For distortion product otoacoustic emissions (DPOAE), we measured the amplitude of evoked otoacoustic emissions to paired pure tones of frequencies f1 and f2, where f1/f2 = 1.2 and the f1 level was 10 dB above f2. Thirty-two sweeps were made in 5 dB steps starting with f1 at 20 dB and ending at 65 dB. The DPOAE threshold was calculated by interpolating the f2 level that would produce a 3 dB response.

### Antibodies

The following primary antibodies were used: mouse IgG1 anti-SIRT3 antibody (1:200; Novus Biologicals; RRID:AB_2818991); goat anti-Oncomodulin antibody (OCM; 1:1000; Santa Cruz; RRID:AB_2267583), rabbit anti-Myosin7a (MYO7; 1:200; Proteus; RRID:AB_10013626) mouse anti-CTBP2 (aka C-Terminal Binding Protein 2; 1:200; BD Transduction Laboratories; RRID:AB_399431), and mouse anti-GRIA2 (aka GluR2/GluA2; 1:2000; Millipore; RRID:AB_2113875). The following secondary antibodies were purchased from Jackson Immuno Research: Donkey Anti-Mouse AF488 (1:500; RRID:AB_2340849), Donkey Anti-Rabbit AF594 (1:500; RRID:AB_2340622), Donkey Anti-Rabbit AF647 (1:200; RRID:AB_2340625), Donkey Anti-Goat AF647 (1:200; RRID:AB_2340438), Goat Anti-Mouse AF594 (IgG1, 1:500; RRID:AB_2338885), AF488 Goat Anti-Mouse (IgG2a, 1:500; RRID:AB_2338855). For the images in Figure 1, an AF568 Goat Anti-Mouse (IgG1, 1:200, Thermo Fisher, RRID: AB_2535766) was used.

**Figure 1.**
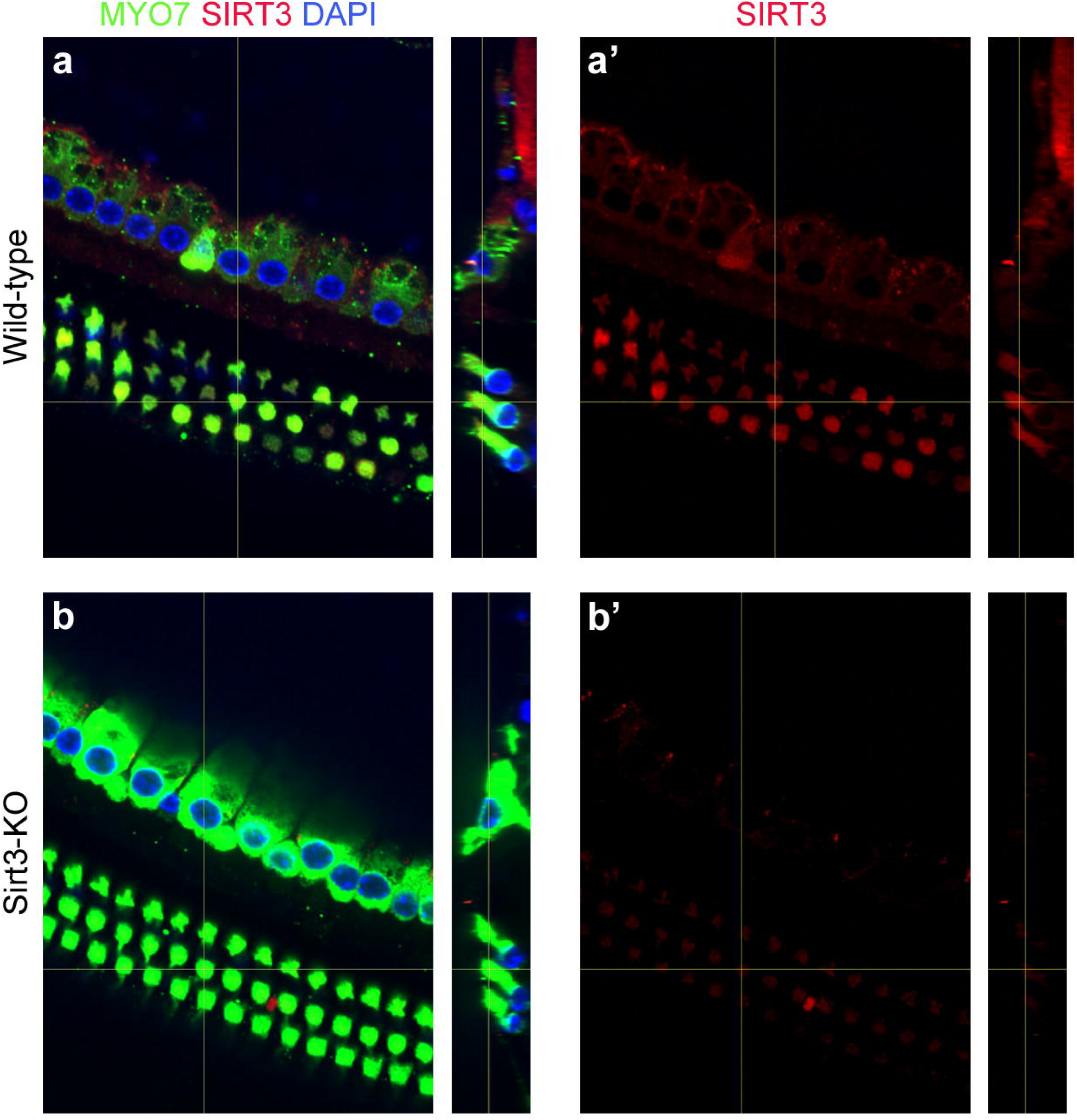
SIRT3 is expressed in sensory cells of the adult mouse cochlea. a. Cochlear whole mount preparation from a wild-type adult mouse, immunolabeled with antibodies specific to MYO7 (green) and SIRT3 (red), and co-stained with DAPI (blue) to reveal nuclei. The vertical yellow lines show the positions from which the side views are taken. (a’) shows anti-SIRT3 immunoreactivity only. b. Cochlear whole mount preparation from a *Sirt3-KO* adult mouse, identically labeled and imaged. This control was used to set the background to reveal SIRT3-specific staining.

### Tissue preparation for immunostaining

Cochlear organs were dissected out of freshly euthanized animals. Their stapes were removed, and a hole was made in their apical tips to allow for adequate fluid exchange. Tissues were immersed in 4% paraformaldehyde (PFA) in PBS for at least overnight, and decalcified in 0.1M EDTA at 4°C on a rotating platform for four days. These cochleae were decalcified in 10% EDTA in 1X PBS (diluted from 10X PBS, Invitrogen) for 2-3 days. All whole mount preparations were microdissected into turns as previously described [26, 27], and the tectorial membrane was carefully removed. Cochleae were mapped using the ImageJ plug-in from Massachusetts Eye and Ear Infirmary.

### Immunostaining

For anti-SIRT3 staining, cochlear pieces were first soaked in 30% sucrose on a shaker for 20 minutes, and then transferred to dry ice for 10-15 minutes or until the sucrose was completely frozen. The samples were allowed to thaw at room temperature and washed with 1X PBS 3 times, rinsing for 20 minutes in between on a shaker. After blocking with 5% serum + 1% Triton, the primary antibody mix diluted in block was added and incubated overnight at 37°C. After rinsing 3 times in PBS (10 minutes each), secondary antibody mix (diluted in block and including Hoechst or DAPI at 1:1000) was added and incubated away from light for 2 hours at 37°C. The pieces were then washed another 3 times with 1X PBS and mounted on slides.

For post-injury histological characterization, dissected mapped turns were immersed in 30% sucrose, flash frozen in liquid nitrogen, allowed to thaw, washed in room temperature Dulbecco’s PBS (Gibco), and blocked for one hour in 1% Triton / 5% donkey serum in PBS. Primary antibody incubations of anti-MYO7, anti-OCM, anti-CTBP2, and anti-GRIA2 were performed at 37°C for 20 hours. The tissue was washed in PBS, and secondary antibody incubation was performed at 37°C for an additional 2 hours, with both MYO7 and OCM labeled with 647-conjugated antibodies. This whole mount protocol was kindly provided by Leslie Liberman. All tissue was mounted using ProLong Gold (Fisher). Whole mounts were placed between two 50 mm coverslips for imaging.

### Confocal microscopy and image processing for figures

Anti-SIRT3 immunostaining was analyzed at the Center for Advanced Microscopy/Nikon Imaging Center at Northwestern University, using a Nikon A1 Laser Scanning or Nikon W1 Dual CAM Spinning Disk imaging setup. Appropriate excitation and emission settings were used and fixed for all panels in the same images presented in the paper. Post-injury histological characterizations were done on an Olympus FV1000 laser scanning confocal microscope at the Center for Advanced Light Microscopy and Nanoscopy at URMC. ImageJ (NIH) was used to Z-project maximal brightness in confocal stacks. Photoshop (Adobe) was used to set maximal and background levels of projections for the construction of figures.

For cochleogram analysis, composite images for cochleograms were assembled in Photoshop by pasting each optical section into its own layer, and merging the pieces of the optical sections where hair cells were evident. Alternatively, projections of confocal stacks were used when individual hair cells could be clearly distinguished. Composite or projected images for the mapped regions of each organ were assembled in a single file in Photoshop. 100 micron lengths were pasted onto the images on the row of pillar cells, and OHCs were counted to determine the percent lost. Where the loss of OHCs was great enough (>30%), an average OHC count, determined from low-frequency regions of the same cochlea, was used as the denominator [28].

Synaptic components of inner hair cells (IHCs) were imaged with a 100X oil lens at 2X magnification. To obtain 3D images of IHCs, the confocal files were opened in Amira 6.0 (Visualization Science Group), appropriately thresholded, and rotated to better display the IHCs. To count ribbon synapses, we visually quantified red/green synaptic puncta, as indicated by colocalized staining within IHCs on projected confocal stacks. The analyst was blinded to genotype, frequency and condition, and the images were randomized in presentation, by renaming the images according to cards drawn from a fair deck. Synapse numbers were then divided by the number of hair cells in the field to obtain a biological replicate. In previous publications, we have used automated synapse detection methods; however, in some of these preparations the staining background for GRIA2 was too variable for computational approaches that use thresholding. This was determined by comparing the number of synapses obtained through visual inspection to those obtained through automated means [29].

### Statistical analysis

Statistical tests for ABR and DPOAE thresholds, and for DPOAE input-output functions, were performed in R using standard functions. The data distribution for cochleograms did not meet the Shapiro test for normality, and was therefore assessed with the Kruskal-Wallis rank sum test. Both functions were also performed in R. To accurately calculate significance levels for amplitude progressions, generalized estimating equations (GEE) were used. These are suitable for the analysis of longitudinal data when population level mean responses suffice to resolve hypotheses. The correlation among responses induced by the repeated measure design were thus modelled by constructing a suitable covariance matrix [30].

Generalized linear model conventions were used. Responses were modeled as Poisson variates, with varying scale parameter to account for overdispersion. The exchangeable correlation model was used to account for within-sample correlation. The link function was *g(μ) = log (μ)* so that *μ e*^*η*^,where *η* is a linear prediction term dependent on threshold x and binary group factor *I.* The null hypothesis of no group effect was modelled as *H*_*0*_ : *η = poly(x, d)*, and the alternative hypothesis was modelled as Hα : *η = poly(x, d)* * *I*, where *poly(x, d)* is an order *d* polynomial with unknown parameters. For the baseline, 14 day post noise (DPN) and KO analyses, *d* = 2 was used, while for the WT analysis *d* = 4 proved to be a significantly better fit. Observed significance levels were obtained both by the χ^2^ approximation method of **geeglm**, as well as by a parametric bootstrap method. P-values obtained by both methods were comparable. However, the bootstrap procedure was the more conservative, so these P-values were used (see Table 1).

**Table 1.**
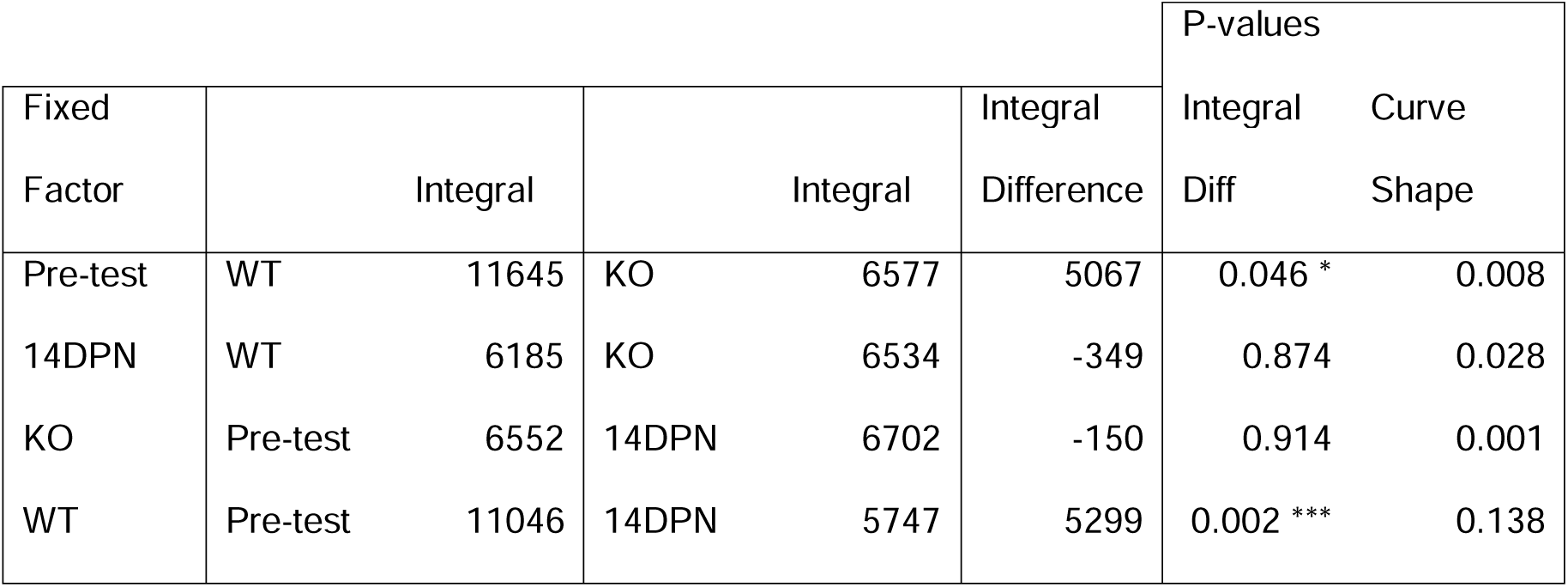
Integrals and p-values for 32 kHz ABR voltage progressions for homozygous *Sirt3-KO* and wild-type mice.

|  |  |  |  |  |  | P-values |  |
| --- | --- | --- | --- | --- | --- | --- | --- |
| Fixed Factor | Integral |  | Integral |  | Integral Difference | Integral Diff | Curve Shape |
| Pre-test | WT | 11645 | KO | 6577 | 5067 | 0.046 * | 0.008 |
| 14DPN | WT | 6185 | KO | 6534 | -349 | 0.874 | 0.028 |
| KO | Pre-test | 6552 | 14DPN | 6702 | -150 | 0.914 | 0.001 |
| WT | Pre-test | 11046 | 14DPN | 5747 | 5299 | 0.002 *** | 0.138 |

## RESULTS

Immunofluorescence for SIRT3 protein reveals its expression pattern in the adult mouse cochlea (Fig. 1). Inner and outer hair cells (IHC & OHC) were visualized with anti-Myosin7 antibodies (Fig. 1, MYO7, green), and nuclei were revealed with DAPI (Fig. 1, blue). SIRT3 immunoreactivity was evident in OHCs and especially in the positions of nerve root endings and supporting cells surrounding the IHCs (Fig. 1a, red). The latter finding is readily observed in the side view (Fig. 1a, cf. red to green). SIRT3 immunoreactivity was significantly greater than that of age-matched, exposure-matched preparations from homozygous *Sirt3-KO* mice (Fig. 1b, red), demonstrating antibody specificity. After multiple attempts, endogenous SIRT3 protein in the cochlea was not sufficiently abundant for us to visualize on a western blot (not shown). Messenger RNA levels for sirtuins have been previously described in the adult mouse cochlea, with *Sirt3* mRNA significantly expressed in adult mouse IHC and OHC [31].

To evaluate SIRT3’s role in hearing recovery, we used a noise dose that confers little permanent damage to awake, two month old, wild-type FVB/nJ mice [22, 24]. An exposure to an 8-16 kHz octave band at 105 dB for thirty minutes is sufficient to elevate auditory brainstem response (ABR) thresholds in mice one day after noise exposure, which is fully recovered in two weeks. This exposure reduces high-frequency peak 1 amplitudes two weeks after exposure, but does not significantly eliminate high-frequency synapses [24]. Mice underwent hearing tests at around P50, followed by noise exposure at P60, a second hearing test at P61, and a third hearing test at P74 (Fig. 2a). At P74, mice were euthanized for histological analysis.

**Figure 2.**
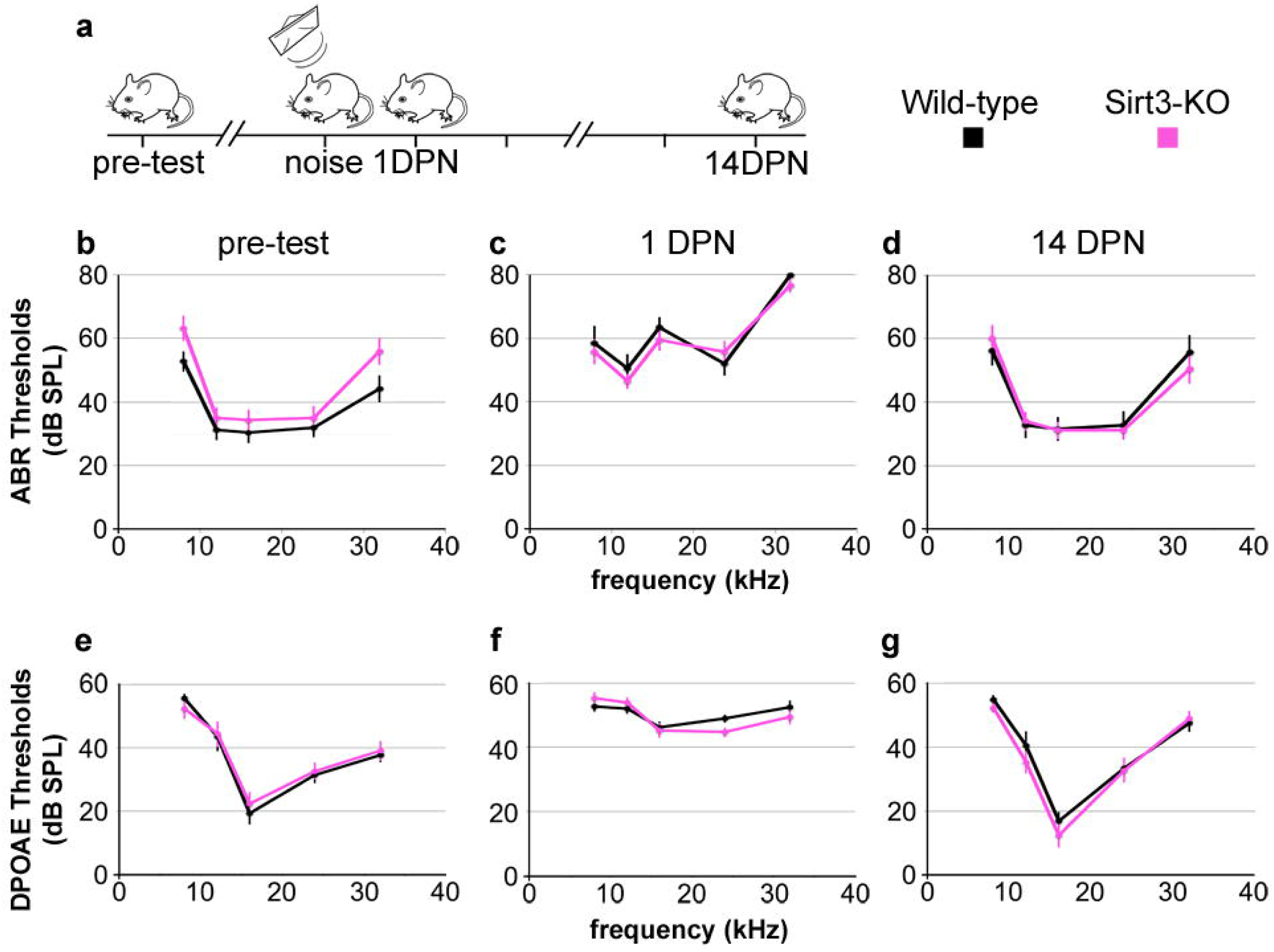
SIRT3 loss of function does not impact endogenous hearing recovery from noise-induced temporary threshold shifts. a. The time course of experiments is shown. Homozygous *Sirt3-KO* mice and wild-type littermates are tested for ABR and DPOAE thresholds prior to noise exposure (pre-test). At P60, they are exposed to an 8-16 octave band noise at 105 dB for 30 minutes (noise), which is sufficient to drive temporary threshold shifts in this strain [22, 24]. Their hearing thresholds are assessed again at 1 day post noise (1 DPN) and 14 days post noise (14 DPN). ABR and DPOAE are assessed at 8, 12, 16, 24, and 32 kHz for all panels. b. Mean ABR thresholds prior to noise exposure for homozygous *Sirt3-KO* mice (n=14, pink) and wild-type littermates (n=13, black). Mice were approximately P50 when tested. Homozygous *Sirt3-KO* mice had significantly worse overall hearing (p=0.004 for genotype, two way ANOVA, n=27 mice total); however, no single frequency was significantly worse (p=0.053, 0.39, 0.41, 0.53, 0.059 for the five frequencies respectively, two-tailed pairwise t-tests at each frequency with Bonferroni correction, n=27 mice total). c. Mean ABR thresholds at 1 DPN for homozygous *Sirt3-KO* mice (n=14, pink) and wild-type littermates (n=13, black). No significant difference was seen between genotypes (p=0.35 for genotype, two-way ANOVA, n=27 mice total). d. Mean ABR thresholds at 14 DPN for homozygous *Sirt3-KO* mice (n=14, pink) and wild-type littermates (n=13, black). No significant difference was seen between genotypes (p=0.85 for genotype, two way ANOVA, n=27 mice total). Overall ANOVA for all three time points showed no significant difference between genotypes (p=0.33, multi-way ANOVA, n=27 mice total). e. Mean DPOAE thresholds prior to noise exposure for homozygous *Sirt3-KO* mice (n=14, pink) and wild-type littermates (n=13, black). No significant difference was seen between genotypes (p=0.43 for genotype, two-way ANOVA, n=27 mice total). f. Mean DPOAE thresholds at 1 DPN for homozygous *Sirt3-KO* mice (n=14, pink) and wild-type littermates (n=13, black). No significant difference was seen between genotypes (p=0.50 for genotype, two-way ANOVA, n=27 mice total). g. Mean DPOAE thresholds at 14 DPN for homozygous *Sirt3-KO* mice (n=14, pink) and wild-type littermates (n=13, black). No significant difference was seen between genotypes (p=0.25 for genotype, two-way ANOVA, n=27 mice total). Error bars: s.e.m.

We found that prior to noise exposure, homozygous *Sirt3-KO* mice had slightly but significantly worse ABR thresholds on average compared to wild-type littermates (Fig. 2b, pink traces, p=0.004 for genotype, two-way ANOVA, n=27 mice total). Notably, no single frequency was significantly worse (p=0.053, 0.39, 0.41, 0.53, 0.059 for 8, 12, 16, 24, and 32 kHz frequencies respectively, two-tailed pairwise t-tests at each frequency with Bonferroni correction, n=27 mice total). At 1 DPN, homozygous *Sirt3-KO* mice had significantly elevated ABR thresholds (Fig 2c, pink traces, p=6 ×10^−14^, for time point, two way ANOVA, n=14 mice). ABR thresholds were also elevated for wild-type littermates (Fig. 2c, black traces, p=2×10^−16^ for time point, two-way ANOVA, n=13 mice). Thresholds for homozygous *Sirt3-KO* and wild-type mice were not different at 1 DPN (Fig. 2c, p=0.35 for genotype at 1 DPN, two-way ANOVA, n=27 mice). At 14 DPN, there was also no difference in ABR thresholds between genotypes (Fig. 2d, p=0.85 for genotype at 14 DPN, two-way ANOVA, n=27). We similarly calculated thresholds for distortion product otoacoustic emissions (DPOAE) for the same mice at the same time points (Fig. 2e-g). DPOAE thresholds for both genotypes were similarly elevated at 1 DPN for both genotypes, and recovered to the same levels at 14 DPN (p=0.13 for genotype, multivariate ANOVA, n=27 mice).

We evaluated the requirement for SIRT3 on ABR peak 1 amplitudes of homozygous *Sirt3-KO* mice and their wild-type littermates before and after noise damage. Mean ABR thresholds for homozygous *Sirt3-KO* mice were 55.7 dB ± 4.1 at 32 kHz, compared to 44.2 ± 4.1 for wild-type littermates, which was not different when adjusted for multiple comparisons. Note that thresholds are determined by the continued presence of any part of the waveform, and not solely by the presence of peak 1. To assess differences between peak 1 amplitude progressions, we used generalized estimating equations to build linear models of the correlated data, as these are suited for the analysis of longitudinal data when population level mean responses suffice to resolve hypotheses [30]. Fig. 3 shows pairwise comparisons of data from each mouse (Fig. 3a-d, light lines), the linear models (Fig. 3a-d, heavy lines) as well as mean latency values at both time points for peak 1 (Fig. e, f). These values of the amplitude integrals and their associated p-values are listed in Table 1 (see Methods for a full description). We propose that the amplitude integrals (Table 1) are appropriate for evaluating differential recruitment of auditory synapses as the stimulus is amplified, when genotypes and time points are compared. Mean peak 1 amplitudes were significantly reduced for homozygous *Sirt3-KO* mice compared to wild-types at 32 kHz, after adjusting for multiple comparison using this approach (p=0.046, parametric bootstrap method, n=27 mice). Any difference between the two datasets was lost at 14 DPN, as mean peak 1 amplitudes at 32 kHz were significantly reduced in wild-type littermates (p=0.002, parametric bootstrap method, n=13 mice) to similar levels as homozygous *Sirt3-KO* mice (p=0.87, parametric bootstrap method, n=27 mice). Curiously, mean peak 1 amplitudes at 32 kHz in homozygous *Sirt3-KO* mice were unchanged between conditions (p=0.914, parametric bootstrap method, n=14). Latencies for peak 1 were not different between genotypes at either time point (Fig. 3e, f, cf. black to red).

**Figure 3.**
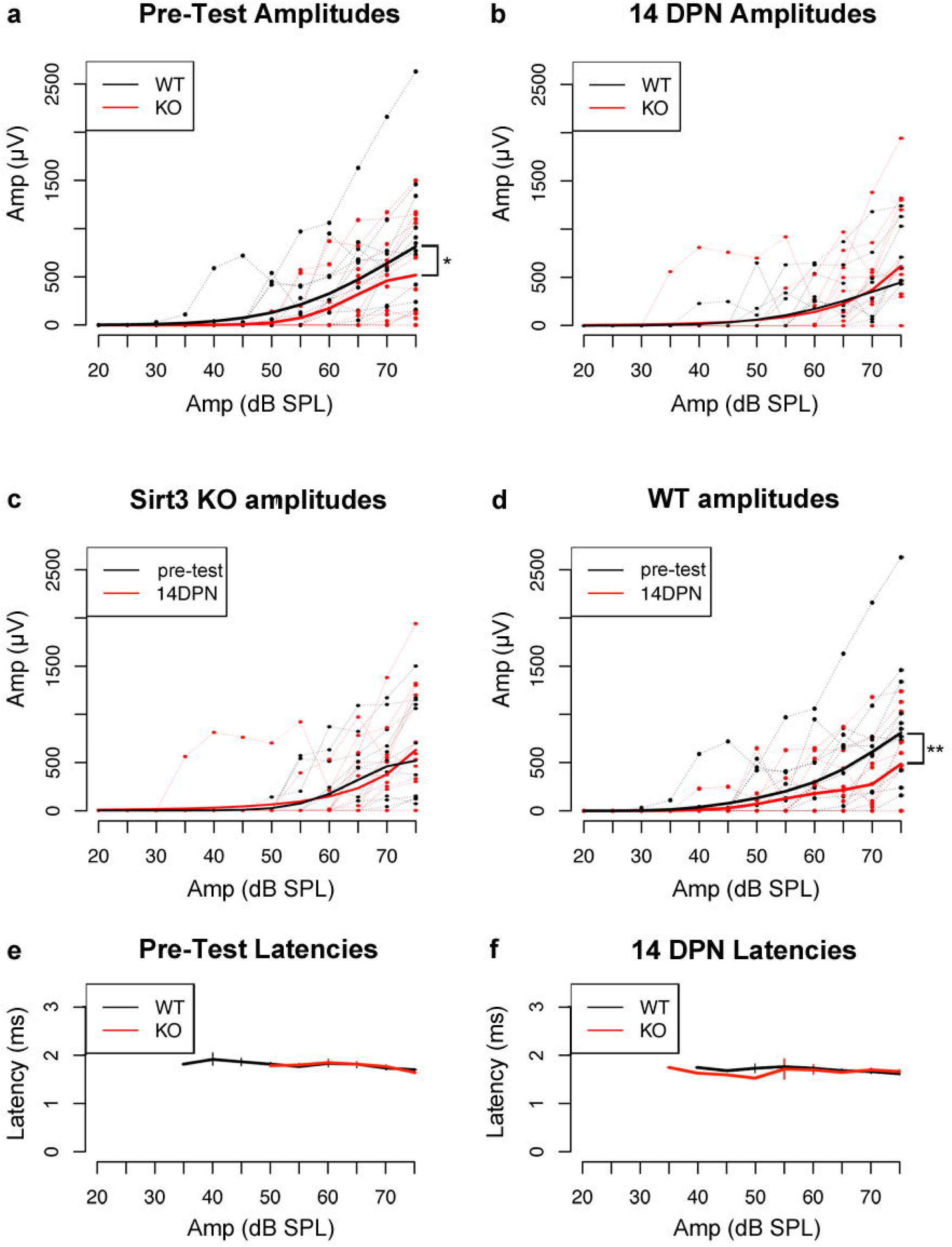
Sirt3 loss of function impacts pre-noise ABR peak 1 amplitudes, but not amplitudes after noise exposure. a. Peak 1 amplitudes in ABR measurements for 32 kHz stimuli prior to noise exposure for homozygous *Sirt3-KO* mice (n=14, thin red) and wild-type littermates (n=13, thin black). Peak 1 amplitudes (microvolts) are plotted on the y axis for different amplitudes of stimulus (dB SPL) plotted on the x-axis. Heavier red and black lines represent respective GEE models. The area under the curve, representative of the progressive neuronal recruitment, is significantly reduced for homozygous *Sirt3-KO* mice (p=0.046, parametric bootstrap method, n=27 mice). b. Same analysis as in (a), but for peak 1 amplitude values obtained from the same mice 14 DPN. No difference is seen between genotypes (p=0.87, parametric bootstrap method, n=27 mice). c. Same analysis as in (a, b), however, in this case amplitudes obtained from homozygous *Sirt3-KO* mice are compared before and after noise exposure. No differences are seen (p=0.914, parametric bootstrap method, n=14). c. Same analysis as in (a, b), however, in this case amplitudes obtained from wild-type littermates are compared before and after noise exposure. A significant reduction in amplitude is evident after noise exposure (cf. red to black, p=0.002, parametric bootstrap method, n=13). e. Mean peak 1 latencies for 32 kHz stimuli prior to noise exposure for homozygous *Sirt3-KO* mice (n=14, pink) and wild-type littermates (n=13, black). No differences are seen between genotypes. f. Mean peak 1 latencies for 32 kHz stimuli at 14 DPN for homozygous *Sirt3-KO* mice (n=14, pink) and wild-type littermates (n=13, black). No differences are seen between genotypes. Error bars: s.e.m.

We evaluated the requirement for SIRT3 on DPOAE fine structure by comparing input-output functions for homozygous *Sirt3-KO* and wild-type mice, prior to noise exposure and at 14 DPN (Fig. 4). Values for each L2 frequency are displayed separately, comparing homozygous *Sirt3-KO* mice (Fig. 4, pink) to wild-type littermates (Fig. 4, black). The curves overlapped at each frequency, and no significant differences were observed (p=0.07 for genotype, multivariate ANOVA, n=27 mice). Taken together, these data suggest that the absence of endogenous SIRT3 activity does not confer susceptibility to permanent functional impairment from noise exposure, by these measures.

**Figure 4.**
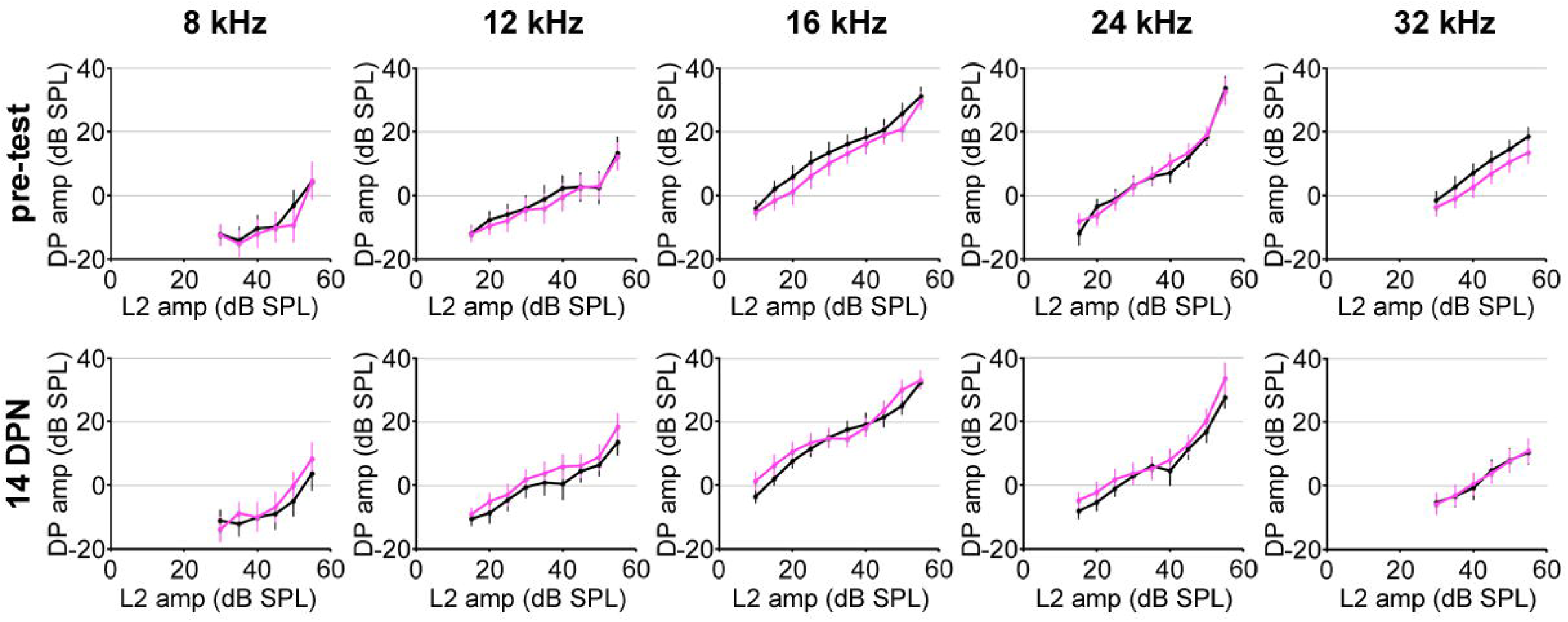
No differences in input-output functions for DPOAE between homozygous Sirt3-KO mice and wild-type littermates at any time point. Ten graphs are arranged by time point, with pre-test results on the top row and 14 DPN results on the bottom row, as well as by frequency, displaying 8 kHz, 12 kHz, 16 kHz, 24 kHz, and 32 kHz results from left to right. Mean distortion product (DP) amplitudes in dB are plotted on the y-axis, and the amplitudes of the L2 stimulus (dB) are plotted on the y-axis. No differences between the genotypes (p=0.07 for genotype, multivariate ANOVA, n=27 mice). Error bars: s.e.m.

Given SIRT3’s immunoreactivity in OHCs (Fig. 1), we sought to determine if these cells are more susceptible to cell death from noise in the absence of SIRT3. We prepared mapped cochlear organs for whole mount imaging of OHCs and IHCs for cochleogram analysis. The cochleae were divided into 100 micron segments along the line of pillar cells, surviving OHCs were quantified for each segment, and the percent of lost OHCs plotted as a function of distance along the cochlear length (Fig. 5). Basal OHCs were more prone to losses in both the wild-type and homozygous *Sirt3-KO* mouse in the absence of noise damage (Fig. 5, cf. b to f), with some variability between biological replicates (Fig. 5b, cf. purple and green). Strikingly, noise exposure did not potentiate OHC losses in the homozygous *Sirt3-KO* mouse (Fig. 5, cf. h to f, p=0.49, Kruskal-Wallis rank sum test, n=3-4 biological replicates). No changes in IHC survival were observed for either genotype in either condition (data not shown). These data indicate that SIRT3 loss of function does not significantly impact OHC survival after mild noise insult.

**Figure 5.**
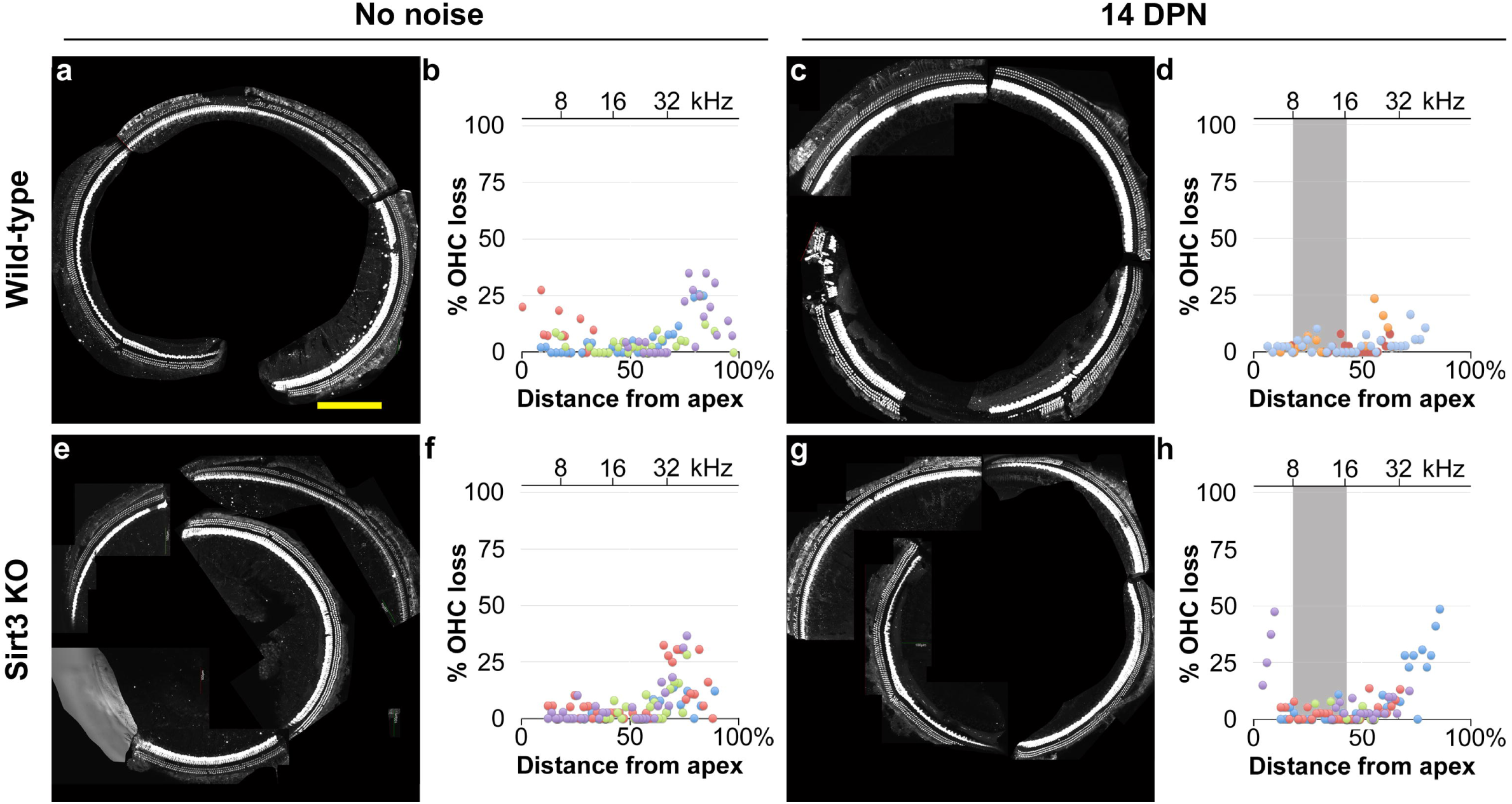
No apparent differences in OHC survival for homozygous *Sirt3-KO* mice and wild-type littermates prior to noise exposure or at 14 DPN. a. Representative cochlear preparation from a wild-type littermate without noise exposure, with IHCs revealed with anti MYO7 antibodies (white) and OHCs revealed with anti-OCM antibodies (also white). Scale bar: 200 microns. b. Cochleogram results from 4 mapped cochlear preparations from 4 wild-type littermates without noise exposure, where OHC loss was quantified in 100 micron segments. The distance from the apex for each segment is plotted on the x-axis, and the percent OHC loss for the segment is plotted on the y-axis. Dots with the same color (purple, blue, green, or red) are from the same cochlear preparation. c. Representative cochlear preparation from a wild-type littermate at 14 DPN, with the same staining as in (a). d. Cochleogram results, similar to (b), from 3 mapped cochlear preparations from 3 wild-type littermates at 14 DPN. Results are similar to (b). e. Representative cochlear preparation from a homozygous *Sirt3-KO* mouse without noise exposure, with the same staining as in (a). f. Cochleogram results, similar to (b), from 4 mapped cochlear preparations from 4 homozygous *Sirt3-KO* mice without noise exposure. Results are similar to (b). g. Representative cochlear preparation from a homozygous *Sirt3-KO* mouse at 14 DPN, with the same staining as in (a). h. Cochleogram results, similar to (b), from 4 mapped cochlear preparations from 4 homozygous *Sirt3-KO* mice at 14 DPN. Results are similar to (f) and (b).

SIRT3 immunoreactivity is also present in cellular structures immediately adjacent to IHCs, around regions of innervation (Fig. 1A). Therefore, we sought to determine if synaptic losses are potentiated after noise exposure in the homozygous *Sirt3-KO* mouse. We imaged MYO7+ IHCs at 12 and 24 kHz that had also been stained with antibodies against CTBP2, the major component of auditory synaptic ribbons, as well as GRIA2, one of the glutamate receptor components of auditory post-synaptic specializations [32]. Representative images for wild-type and homozygous *Sirt3-KO* cochleae, with and without noise exposure, are depicted in Fig. 6. Confocal stacks were imported into Amira, so that the IHCs could be rotated, to view at similar angles. Paired synaptic structures were quantified visually on randomized projected images, whose condition and genotype were masked from the analyst. Each frequency, condition, and genotype is labeled with the mean number of synapses per IHC for 3-4 biological replicates. Wild-type cochlea harbored between 16.7 and 18.0 synapses per IHC in no noise condition (Fig. 6a, b). These numbers were not significantly reduced after noise exposure (Fig. 6c, d), similar to previous results [22, 24]. At both frequencies and in both conditions, homozygous *Sirt3-KO* cochleae harbored between 17.8 and 18.8 synapses per IHC (Fig. 6i-l). Thus, we find no evidence that the loss of SIRT3 protein potentiates synaptopathy from noise exposure.

**Figure 6.**
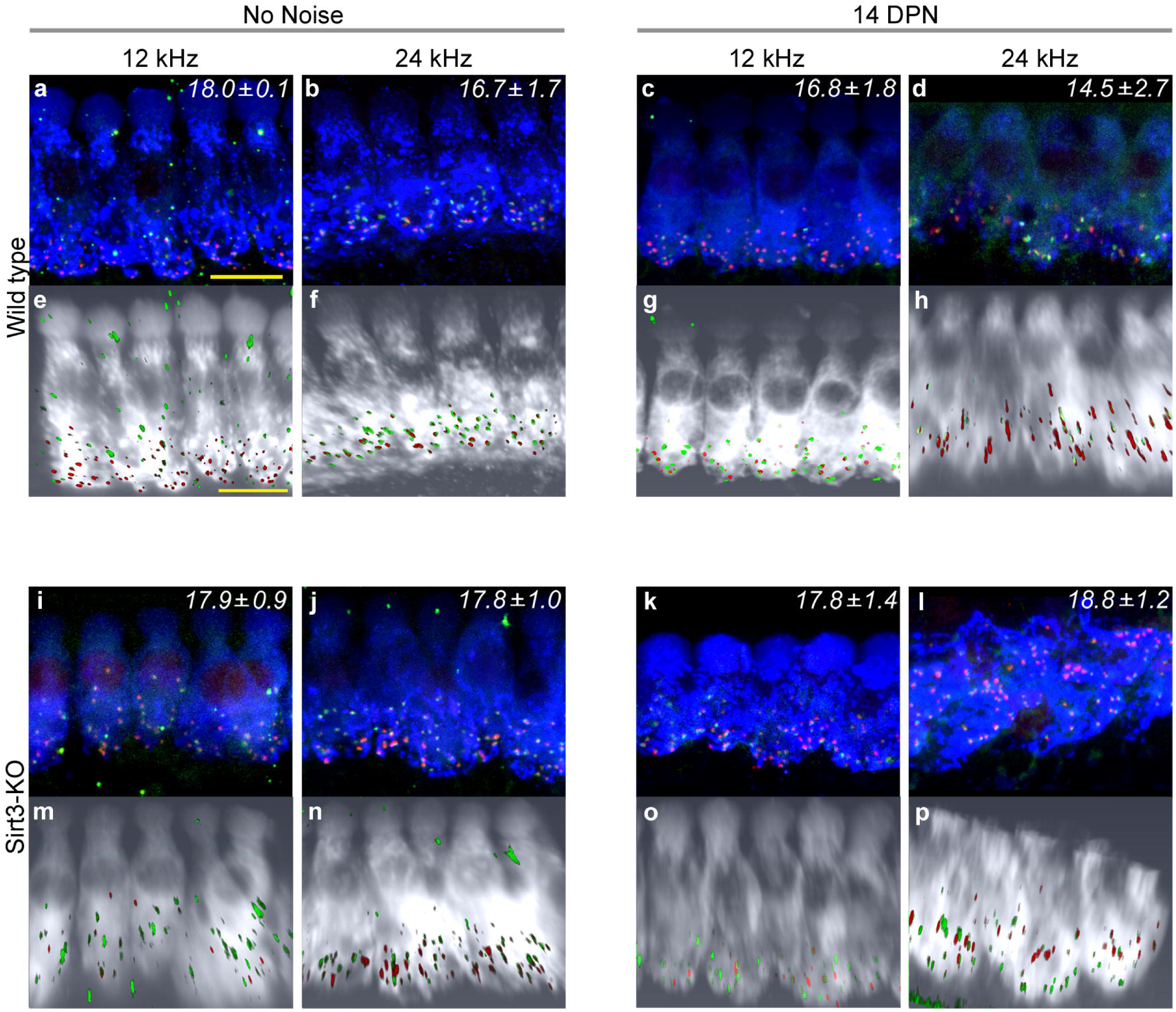
Synaptic loss is not potentiated in homozygous *Sirt3-KO* mice after noise exposure. a. Representative image of wild-type 12 kHz IHCs with no noise exposure, stained with anti-MYO7 (blue), anti-CTBP2 (red) to reveal pre-synaptic structures, and anti-GRIA2 (green) to reveal post-synaptic structures. Mean synaptic number per IHC (± s.e.m.), visually counted from 3 biological replicates with 8 IHCs each, is noted in the upper right hand corner. Size bar: 5 microns. b. Representative image of wild-type 24 kHz IHCs with no noise exposure, similarly stained to (a). 3 replicates, 6-7 IHCs each. c. Representative image of wild-type 12 kHz IHCs at 14 DPN, similarly stained to (a). 4 replicates, 7-11 IHCs each. d. Representative image of wild-type 24 kHz IHCs at 14 DPN, similarly stained to (a). 3 replicates, 7-8 IHCs each. e – h. Three dimensional Amira rendering of confocal stacks shown in (a – d), respectively, with MYO7 rendered in white, CTBP2 in red, and GRIA2 in green. Hair cells are rotated to display from similar orientations. i. Representative image of homozygous *Sirt3-KO* 12 kHz IHCs with no noise exposure, similarly stained to (a). 3 replicates, 6-8 IHCs each. j. Representative image of homozygous *Sirt3-KO* 24 kHz IHCs with no noise exposure, similarly stained to (a). 3 replicates, 4-7 IHCs each. k. Representative image of homozygous *Sirt3-KO* 12 kHz IHCs at 14 DPN, similarly stained to (a). The angle of the tissue in the imaging foreshortens the IHC. 5 replicates, 6-8 IHCs each. l. Representative image of homozygous *Sirt3-KO* 24 kHz IHCs at 14 DPN, similarly stained to (a). IHCs appear different because of the angle of the tissue. 3 replicates, 6-8 IHCs each. m - p. Three-dimensional Amira rendering of confocal stacks shown in (i – l), respectively, with MYO7 rendered in white, CTBP2 in red, and GRIA2 in green. Hair cells are rotated to display from similar orientations. The apical truncation shown in (p) is an artifact of the confocal imaging.

## DISCUSSION

NIHL is a life-long, progressive disability that presents a significant social and personal burden. Variants in genes that reduce oxidative stress can enhance susceptibility to occupational NIHL [7], indicating the importance of this pathway to acquired hearing disorders. SIRT3 is a major regulator of the mitochondrial oxidative stress response [33]. We tested the hypothesis that loss of SIRT3 function could impact endogenous recovery from a mild noise insult in young adult mice. We show that SIRT3 immunoreactivity is present in OHCs, around the post-synaptic structures of IHCs, and faintly within IHCs themselves, comparable to mRNA studies [31]. We measured auditory thresholds of homozygous *Sirt3*-KO mice and their wild-type littermates with ABR and DPOAE, before noise exposure, immediately after noise exposure, and a third time after the mice had recovered. No differences were observed between the genotypes after noise exposure. We did not observe permanent adverse effects on homozygous *Sirt3-KO* OHCs, either by DPOAE fine structure analysis or in their survival after noise damage. We also did not observe any potentiation of high-frequency synaptic loss in homozygous *Sirt3-KO* mice. We did observe a slight but significant reduction in peak 1 amplitude at 32 kHz for homozygous *Sirt3-KO* mice, but this effect was evident prior to noise damage and was not further potentiated. In this finding, we present a new statistical analysis for peak 1 comparisons. Taken together, these results strongly indicate that SIRT3 is not required in mice for endogenous recovery from a single noise exposure.

The oxidative stress response acts to mitigate the effects of traumatic noise injury on the cochlea [11]. Mitochondrial genetic defects are strongly associated with NIHL susceptibility [10], and strategies to reduce mitochondrial oxidative stress can protect from NIHL in animal models [34, 35]. SIRT3 protects by activating mitochondrial enzymes that reduce oxidative radicals, including SOD2 [36] and Catalase [37], through lysine deacetylation. Notably, the exogenous activation of SIRT3 through pharmacological means restores partial cochlear function after traumatic noise damage [20] and aminoglycoside damage [38]. In analyses of SIRT3 effector molecules, heterozygous *Sod2-KO* mice were shown to incur more damage from traumatic noise compared to wild-type controls, including higher ABR thresholds and greater OHC losses [39]. Together, these published data show that in mice, the mitochondrial oxidative stress response is activated after traumatic noise, reduced levels of those effectors worsen trauma outcomes, and increasing the stress response through greater levels of SIRT3 activity improves trauma outcomes. This is consistent with current models of human occupational NIHL susceptibility, and informs some efforts to combat NIHL [40, 41].

In contrast, the experiments in this report assess endogenous recovery after a single sub-traumatic noise exposure. We have previously shown that this paradigm reveals susceptibility to noise damage in homozygous *Foxo3-KO* mice [22]. Others have used similar paradigms to investigate the protective roles of the medial olivocochlear system [42] and of peroxisomes [43]. In all three examples, a sub-traumatic noise exposure readily revealed a genetic susceptibility to damage. Given SIRT3’s expression in OHCs (Fig. 1), its role in activating the oxidative stress response, and the sheer importance of that response in mitigating traumatic noise damage, we were surprised to find no sub-traumatic susceptibility with SIRT3 loss-of-function mice (Figs. 2, 4, & 5). These data may be interpreted to indicate that for a single, sub-traumatic noise exposure, some other cellular protective mechanism is sufficient for endogenous recovery. Heat shock proteins [44] and the FOXO3-dependent response [22] are candidates. Alternatively, a clue may lie in the expression pattern of SIRT3: it may be that the oxidative stress response in SIRT3-negative cells, including supporting cells and/or the stria vascularis, is sufficient to protect OHCs from sub-traumatic noise. We emphasize that these results indicate that cochlear responses to chronic insults, such as aging and occupational noise exposure, contain different or additional components, when compared to cochlear responses to a single, acute insult. In fact, a profound relationship between TTS and age-related hearing loss has been reported, in which acute synaptopathy was associated with an exacerbated long-term change of cochlear function [21]. In this regard, whether the presence of SIRT3 makes a difference in the situation of cochlear synaptopathy or in long-term changes after sub-traumatic noise exposure is worth further investigation.

We also show that high-frequency cochlear synaptopathy from noise exposure is not potentiated in the homozygous *Sirt3-KO* mouse (Fig. 6). This result was also counterintuitive, given the prominent role of excitotoxic glutamate in auditory synaptic loss [21, 45-47] and the presence of SIRT3 immunoreactivity around the synaptic region of the IHCs (Fig. 1). The latter finding suggests that SIRT3 may promote mitochondrial function in neurites and supporting cells, and would be predicted to be protective. However, we note that the homozygous *Sirt3-KO* mouse would have had mitochondrial dysfunction throughout its development. It is possible that compensatory mechanisms to prevent calcium overload are present in homozygous *Sirt3-KO* neurites. Such mechanisms could take the form of changes in efferent neurotransmission. Exogenous dopamine, for example, can lower cochlear action potential amplitudes in guinea pigs, providing a tonic effect that counteracts excitotoxicity [48]. Alternatively, stress signaling within spiral ganglion afferent neurites could modulate their activity. Altered cAMP levels within spiral ganglion neurites, for example, modulate an inward potassium current through HCN channels, which shapes EPSP waveforms and modulates the probability of spike formation [49]. Other explanations are also possible.

The amplitude progressions of ABR peak 1 represent the only significant difference observed between homozygous *Sirt3-KO* mice and their wild-type littermates prior to noise exposure (Fig. 3). We present the novel use of an advanced statistical method, generalized estimating equations, to more accurately calculate significant differences between amplitude progressions [30]. While the amplitude progression is sometimes modeled as a straight line, we contend that this is an unnecessary oversimplification, given the newly described molecular diversity of the responding spiral ganglion population [50]. Generalized estimating equations can provide a function that is a more accurate representation of the progression. Any increases seen in the amplitude progression reflect additional spiral ganglion neurons participating in the synchronized response, as the tone pip gets louder. Since the summation of neuronal activity is the variable of interest, we contend that the integral of the amplitude progression function is the appropriate measure for comparison. Our analysis contrasts with comparing the slopes of two progressions, as these are the derivatives of the neuronal summation. Of course, this method is only valid when thresholds are not statistically different, as we had found for the 32 kHz frequency (Fig. 2). It is also important to point out that the individual scoring the threshold must score on the whole waveform and not solely on the presence or absence of peak 1.

Compensatory mechanisms could also account for the reduced peak 1 amplitude observed in homozygous *Sirt3-KO* mice prior to noise exposure (Fig. 3). Note that in the wild-type mice, peak 1 amplitudes are significantly reduced in response to noise, as we have previously reported [24]. These differences in peak 1 amplitude are not due to synaptopathy, as the numbers of synapses are not different between the genotypes or between conditions. Instead, it must be due to either reduced or desynchronized firing by spiral ganglion neurons. These findings underscore the adaptive nature of the adult cochlea in its response to stimuli in the context of genomic variability, and indicate that cellular stress responses additional to the oxidative stress response promote cochlear homeostasis for sub-traumatic challenges.

## ACKNOWLEDGEMENTS

This work was supported by the NIDCD with R01 DC014261, awarded to PMW, and in part by a grant from the American Hearing Research Foundation, awarded to XT. The confocal imaging for SIRT3 expression was performed at the Northwestern University Center for Advanced Microscopy, generously supported by NCI CCSG P30 CA060553 awarded to the Robert H Lurie Comprehensive Cancer Center. We gratefully acknowledge Dr. Anne Luebke, who maintains the Small Animal Auditory Testing Core and Dr. V. Kaye Thomas, who maintains the Center for Advanced Light Microscopy and Nanoscopy, both at URMC. We also thank Ms. Holly Beaulac for a critical reading of the manuscript.

SP, XT, and PMW planned experiments; SP, LS, ND, and XT performed experiments; LS, ND, XT, AA, and PMW analyzed data; PMW wrote the manuscript draft; and XT, AA, and PMW edited the manuscript. The authors have no competing interests.

